# Wnt7a is Required for Regeneration of Dystrophic Skeletal Muscle

**DOI:** 10.1101/2024.01.24.577041

**Authors:** Uxia Gurriaran-Rodriguez, Kasun Kodippili, David Datzkiw, Ehsan Javandoost, Fan Xiao, Maria Teresa Rejas, Michael A. Rudnicki

## Abstract

Intramuscular injection of Wnt7a has been shown to accelerate and augment skeletal muscle regeneration and to ameliorate dystrophic progression in *mdx* muscle, a model for Duchenne muscular dystrophy (DMD). However, loss-of-function studies to investigate the requirement for Wnt7a in muscle regeneration has not been evaluated. Here, we assessed muscle regeneration and function in wild type (WT) and *mdx* mice where Wnt7a was specifically deleted in muscle using a conditional *Wnt7a* floxed allele and a *Myf5-Cre* driver. We found that both WT and *mdx* mice with deletion of Wnt7a in muscle, exhibited marked deficiencies in muscle regeneration at 21 d following cardiotoxin (CTX) induced injury. Unlike WT, deletion of Wnt7a in *mdx* resulted in a marked decrease in specific force generation prior to CTX injury. However, both WT and *mdx* muscle lacking Wnt7a displayed decreased specific force generation following CTX injection. Notably the regeneration deficit observed in *mdx* mice lacking Wnt7a in muscle was rescued by a single tail vein injection of an extracellular vesicle preparation containing Wnt7a (Wnt7a-EVs). Therefore, we conclude that the regenerative capacity of muscle in *mdx* mice is due to the upregulation of endogenous Wnt7a following injury, and that systemic delivery of Wnt7a-EVs represents a therapeutic strategy for treating DMD.

## Introduction

Wnt7a signaling stimulates muscle regeneration and represents an important intrinsic mechanism for regenerative myogenesis (1, 2). Wnt7a signaling via the Frizzled7 (Fzd7) receptor activates the planar-cell-polarity (PCP) pathway to drive symmetric expansion of muscle stem cells (MuSCs) and accelerates and augments muscle regeneration (2). Wnt7a/Fzd7 signaling in myofibers directly activates the Akt/mTOR growth pathway to induce hypertrophy (3). Wnt7a/Fzd7 also acts to increase the directional migration of satellite cells and progenitors through activation of the PCP effector proteins Dvl2 and the small GTPase Rac1 (1).

Duchenne muscular dystrophy (DMD) is a devastating muscle wasting disease caused by loss-of-function mutations in the dystrophin gene. Loss of dystrophin in myofibers results contraction-induced damage, increased sarcolemma permeability, cytosolic calcium overloading, oxidative and nitrosamine stress, cell dysfunction, and cell death. Dystrophin is also expressed in MuSCs where it functions to establish polarity which is necessary for efficient asymmetric divisions, and generation of progenitors (4). Compromised muscle regeneration combined with ongoing muscle damage leads to the progressive disease phenotype in DMD.

Wnt7a has been demonstrated to have therapeutic potential as a treatment for DMD (5). Intramuscular injection (IM) of recombinant Wnt7a protein into the *tibialis anterior* (TA) muscle in *mdx* mice, a mouse model of DMD, results in an increase in muscle mass, and in myofiber hypertrophy, resulting in a doubling in force generation and a 6-fold reduction in myofiber necrosis (5). These findings suggested that Wnt7a is a promising candidate as an ameliorative therapy for patients with DMD. However, a significant issue in the clinical development of Wnt7a as a therapeutic is that Wnt proteins are highly hydrophobic and thus cannot delivered via the circulation (6).

We have recently reported that Wnt7a is highly expressed by regenerating myofibers following an acute injury induced by cardiotoxin (CTX) injection where Wnt7a is secreted on the surface of exosomes, a class of small extracellular vesicles (EVs) (7). Small EVs carry a variety of cargo and thus facilitate the transfer of proteins (e.g. Wnt, HH, Notch), mRNAs, and microRNAs between cells and tissues (8–10). EVs have been shown to carry Wnts such as Wnt1, Wnt3a, Wnt5a, and Wnt7a on their surface, and efficiently induce Wnt signaling in target cells (7, 11–13), facilitating long-range intercellular communications (14). Systemic injection of EVs has successfully delivered microRNAs and siRNAs to specific sites, such as the brain or tumors, where gene modulation was documented (15–19). Therefore, systemic delivery of Wnt7a on EVs represents a potential strategy for stimulating muscle repair in DMD.

In *mdx* mice, muscle regeneration following an acute injury induced by CTX injection results in relatively efficient regeneration associated with an improvement in the muscle phenotype (20). We hypothesized that the relatively efficient regeneration is due to the high-level induction of endogenous Wnt7a in newly differentiated myofibers that occurs following CTX injury. Therefore, we assessed regeneration following acute injury induced by CTX injection in *mdx* carrying a conditional Wnt7a allele.

We observed a striking deficit in muscle regeneration in mice lacking Wnt7a resulting in reduced muscle function. Similarly to wild type mice, we found that Wnt7a is also secreted at high levels in *mdx* mice in the newly formed regenerating myofibers that arise following an acute muscle injury. Importantly, the deficiency in muscle regeneration in *mdx* muscle lacking Wnt7a could be rescued by a single tail vein injection of a preparation of EVs carrying Wnt7a. Our results indicate that the relatively efficient regeneration observed after acute injury of muscle in *mdx* mice is therefore a consequence of high-level expression of endogenous Wnt7a during regenerative myogenesis. Moreover, our experiments suggest that systemic delivery of Wnt7a-EVs via the circulation represents a potential therapeutic approach for treating DMD.

## Results

### Wnt7a is required for efficient muscle regeneration in *mdx* mice

We hypothesized that the relatively efficient regeneration in *mdx* muscle following cardiotoxin (CTX)-induced injury is due to the high-level induction of endogenous Wnt7a that occurs in newly differentiated myofibers (2). Accordingly, we observed high level induction of Wnt7a in newly differentiating myofibers in the *tibialis anterior* (TA) muscles in *mdx* mice at 96 h following CTX injury (Fig. 1A). Therefore, we evaluated regenerating muscle from mice with a functional Wnt7a gene (*Myf5^Cre/+^:Wnt7a^+/+^*) versus mice where Wnt7a was specifically deleted in muscle (*Myf5^Cre/+^:Wnt7a^fl/fl^*), in both *C57/BL10* (WT) or *C57BL/10ScSn-Dmd^mdx^/J* (*mdx*) backgrounds (Fig. 1B).

**Figure 1:**
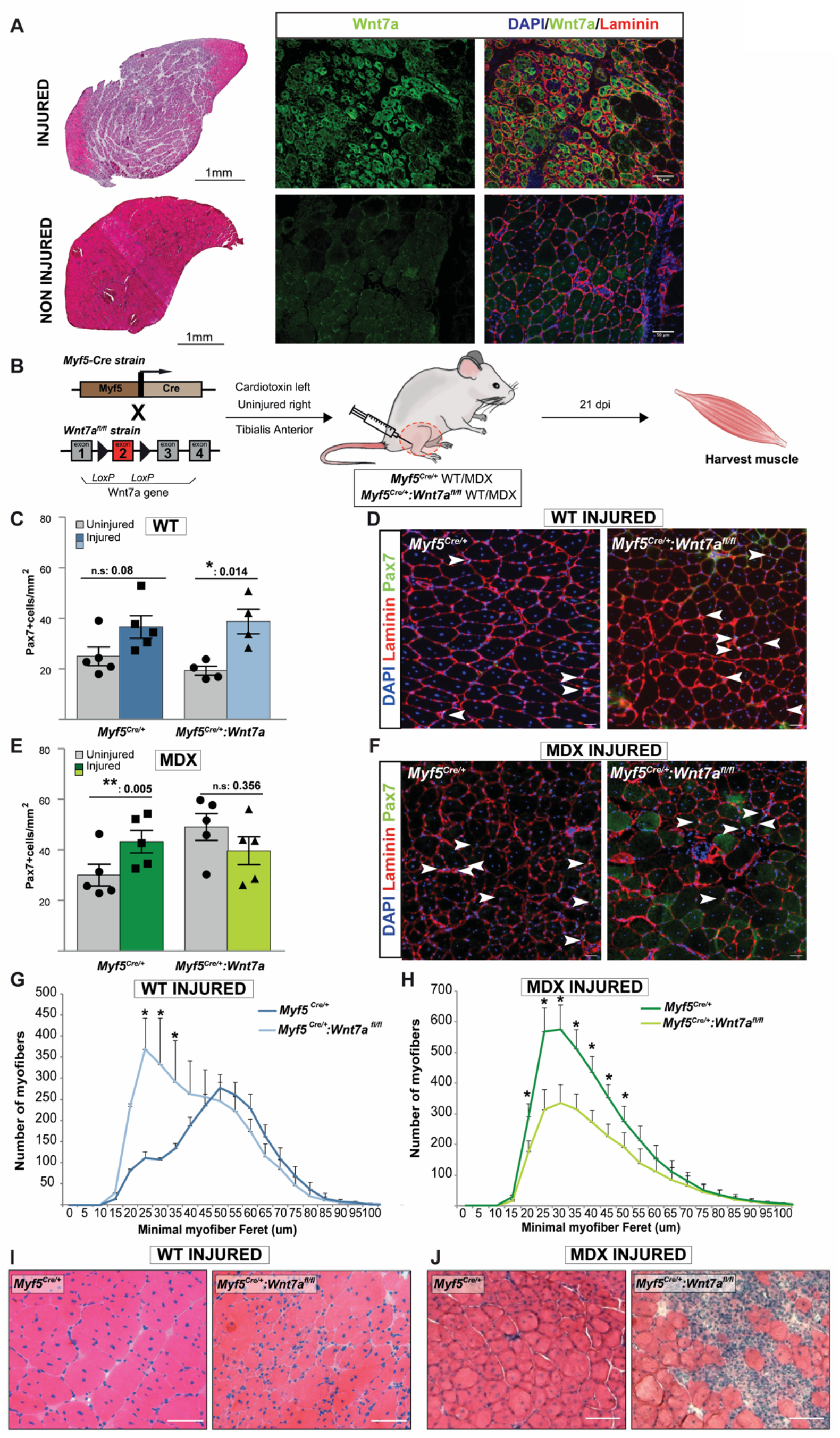
Lack of Wnt7a expression impairs muscle regeneration. (*A-left*) H&E stained sections of injured and non-injured TA muscle of *mdx* mice at 3 dpi. (*A-right*) Wnt7a immunostained (green) sections of injured and non-injured TA upon 3 dpi. Laminin (red) delineates myofibers. Nuclei are stained with DAPI. (*B*) Mouse strains used and experimental design. (*C* and *D*) Quantification of Pax7-expressing cells in regenerating WT muscle. (*E* and *F*) Quantification of Pax7-expressing cells in regenerating *mdx* TA muscle. (*G* and *H*) Myofiber caliber distribution at 21 d following regeneration in WT and *mdx* TA muscle with and without Wnt7a deletion. (*I* and *J*) H&E stained sections at 21 d following regeneration in WT and *mdx* TA muscle with and without Wnt7a deletion. TA (Tibialis Anterior), dpi (days post injury). n≥4 mice, mean ± s.e.m., p values determined by two-sided Student’s t-test (*p<0.05, n.s.=not significant).

In uninjured TA muscles from WT mice, no significant difference was observed in the numbers of Pax7-expressing cells regardless of whether Wnt7a was deleted (Fig. 1C). At 21 days following CTX injury (Fig. 1C, 1D), both strains exhibited about a 2-fold increase in the numbers of Pax7-expressing cells. In WT mice lacking Wnt7a, analysis of minimal fiber Feret in the TA at 21d post-CTX revealed a significant shift towards smaller myofibers and a corresponding reduction in larger myofibers (Fig. 1G-I and S1A). We also observed a significant increase in the total number of myofibers (Fig. S1B). However, no significant difference was observed in the TA total cross-sectional area (Fig. S1C). Therefore, we conclude that Wnt7a is required for normal progression of muscle regeneration in WT mice.

In uninjured TA muscles from *mdx* mice, we observed about a 2-fold increase in numbers of Pax7-expressing cells when Wnt7a was deleted (Fig. 1E). In contrast to WT mice, deletion of Wnt7a in *mdx* muscle did not result in an increase in numbers of Pax7-expressing satellite cells at 21 d following CTX injection (Fig. 1E, 1F). Interestingly, examination of TA muscle at 21d following CTX-induced injury revealed a similar distribution in myofiber ferret sizes (Fig. 1H), and no significant differences in the average myofiber caliber (Fig. S1D). However, we noted significantly decreased numbers of myofibers present in the regenerated *mdx* muscle lacking Wnt7a (Fig. S1E), but no difference in TA total cross-sectional area (Fig. S1F). Finally, we observed extensive areas of fibrosis that were not evident in regenerated muscles in *mdx* mice with Wnt7a (Fig. 1J).

Overall, these data are consistent with an essential role played by Wnt7a/Fzd7 signaling in stimulating the efficient regeneration of skeletal muscle after an acute injury. Our experiments indicate that in the absence of Wnt7a, the regeneration deficit in dystrophin deficient muscle is exasperated resulting in a significant impairment in the regenerative response.

### Reduced force generation in regenerated muscles lacking Wnt7a

To evaluate the functional consequence of a lack of Wnt7a on muscle regeneration, we examined force generation in WT and *mdx* mice that either expressed Wnt7a (*Myf5^Cre/+^:Wnt7a^+/+^*), or where Wnt7a had been deleted (*Myf5^Cre/+^:Wnt7a^fl/fl^*) (Fig. 2A). In uninjured TA muscles from WT mice, deletion of Wnt7a in muscle did not have a significant effect on specific force generation across the force frequency profile (Fig. 2B). By contrast, in uninjured TA muscles from *mdx* mice lacking Wnt7a, we observed a significant decrease in specific force generation relative to uninjured TA muscles from *mdx* mice expressing endogenous Wnt7a (Fig. 2C).

**Figure 2:**
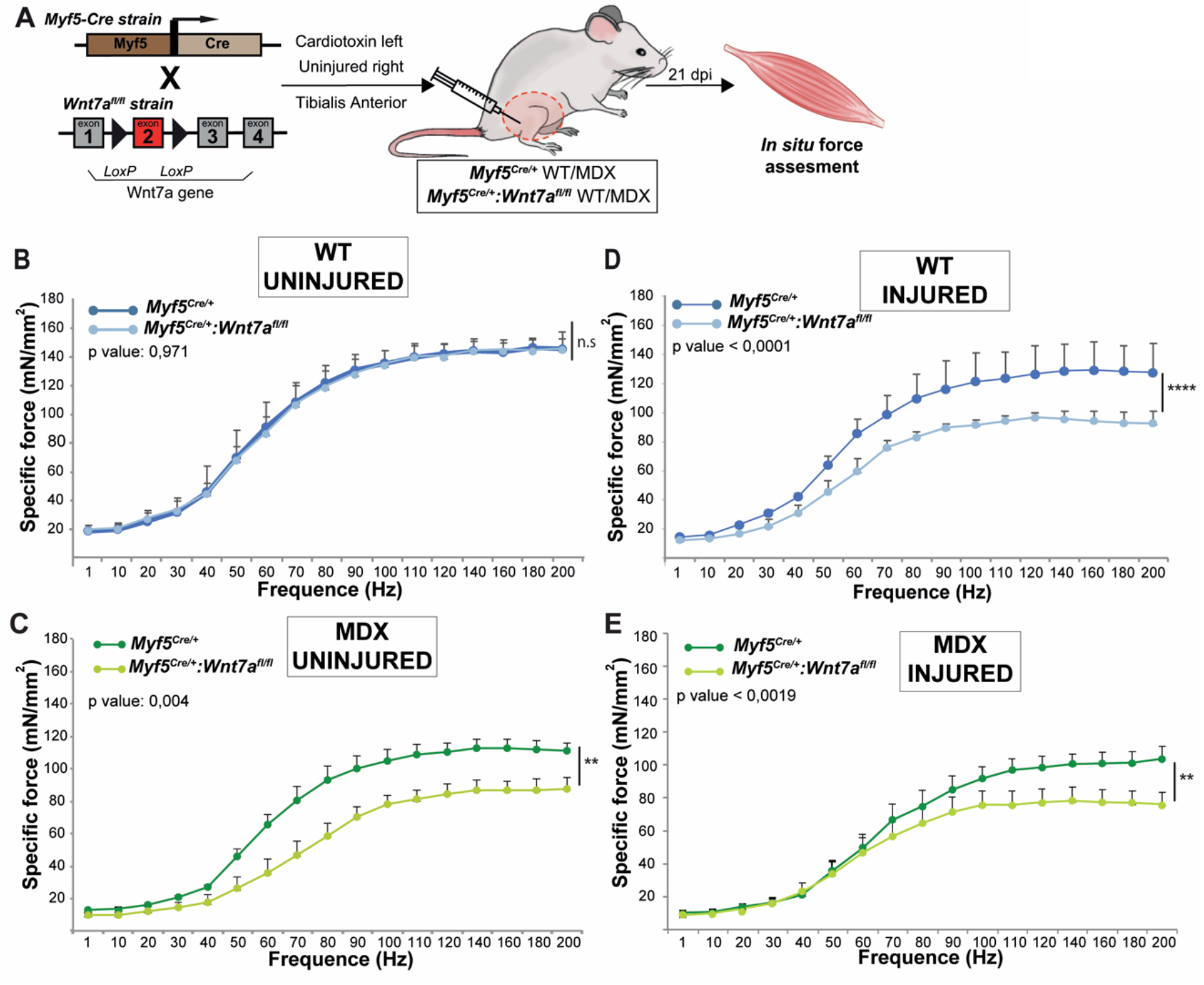
Reduced force generation in regenerated TA muscles lacking Wnt7a. (*A*) Schematic of mouse strains used and experimental design. (*B*) Specific force frequency curve in uninjured WT TA muscle with (dark blue) and without (light blue) Wnt7a expression. (*C*) Specific force frequency curve in uninjured *mdx* TA muscle with (dark green) and without (light green) Wnt7a expression. (*D*) Specific force frequency curve of regenerated WT TA muscle, with (dark blue) and without (light blue) Wnt7a expression, at 21 d following injury. (*E*) Specific force frequency curve of regenerated *mdx* TA muscle, with (dark green) and without (light green) Wnt7a expression, at 21 d following injury. TA (Tibialis Anterior). n≥3 mice, mean ± s.e.m., p values determined by two-way ANOVA test (*p<0.05, **p<0.01, n.s.=not significant).

Examination of force generation in the regenerated TA muscle at 21 d following CTX injury revealed a significant reduction in specific force generation in WT mice lacking Wnt7a (Fig. 2D). This was associated with a significant increase in the weight of the CTX injured TA (Fig. S2A, S2B). These results support the notion that Wnt7a is required for the efficient regeneration of muscle following an acute injury. In *mdx* TA muscle lacking Wnt7a, we observed a significant reduction in specific force generation at 21 d after CTX injection (Fig. 2E). In contrast to WT muscle, no significant change in the weight of the CTX injured TA was observed (Fig. S2C, S2D). Together, these results support the conclusion that Wnt7a is required for the normal regenerative response following an acute muscle injury in both WT and mdx muscle. Furthermore, the reduced force generation in uninjured TA muscles of *mdx* mice lacking Wnt7a argues that Wnt7a is playing a role in slowing the progression of dystrophic disease due to chronic muscle damage in *mdx* mice.

### Wnt7a on extracellular vesicles display bioactivity

We were interested in knowing whether Wnt7a delivered systemically loaded on EVs to *mdx* mice, where Wnt7a was deleted in muscle, would rescue the regeneration deficit. To assess the bioactivity of Wnt7a-EVs on MuSCs, we first asked whether Wnt7a-EVs could stimulate symmetric MuSCs division as previously described using recombinant Wnt7a(2). Therefore, EVs were isolated from Wnt7a (Wnt7a-EV) and empty vector (CTR-EV) transfected primary myoblasts after three days of differentiation using a tangential flow filtration (TFF) protocol we developed (7) (Fig. S3A). Myotube derived small EVs exhibited the typical size range of between 50-200nm (Fig. S3B). Examination of purified Wnt7a-EVs by immunogold transmission electron microscopy (iTEM) after staining using anti-Wnt7a antibody revealed localization of Wnt7a to the surface of EVs, with the appropriate size and morphology (Fig. S3C). Wnt7a protein content on isolated EVs was quantified using a standard curve of Wnt7a recombinant protein and CD63 as a marker of EVs (Fig. S3D). We observed a content of 2 ng of Wnt7a per μg of total EV protein content.

Myofibers from the *Flexor Digitorum Brevis* (FDB) muscles were isolated from *Myf5^Cre/+^:R26R-eYFP* mdx mice (Fig. 3A) and cultured in 96-well plates and high content screening performed as previously described (21). Isolated FDB fibers were treated with 20 ng/ml of Wnt7a equivalents delivered on EVs (Wnt7a-EV), alternatively with recombinant protein (rWnt7a), or also CTR-EVs. FDB myofibers were cultured for 42h, at which point the majority of muscle stem cells have undergone one round of cell division.

**Figure 3:**
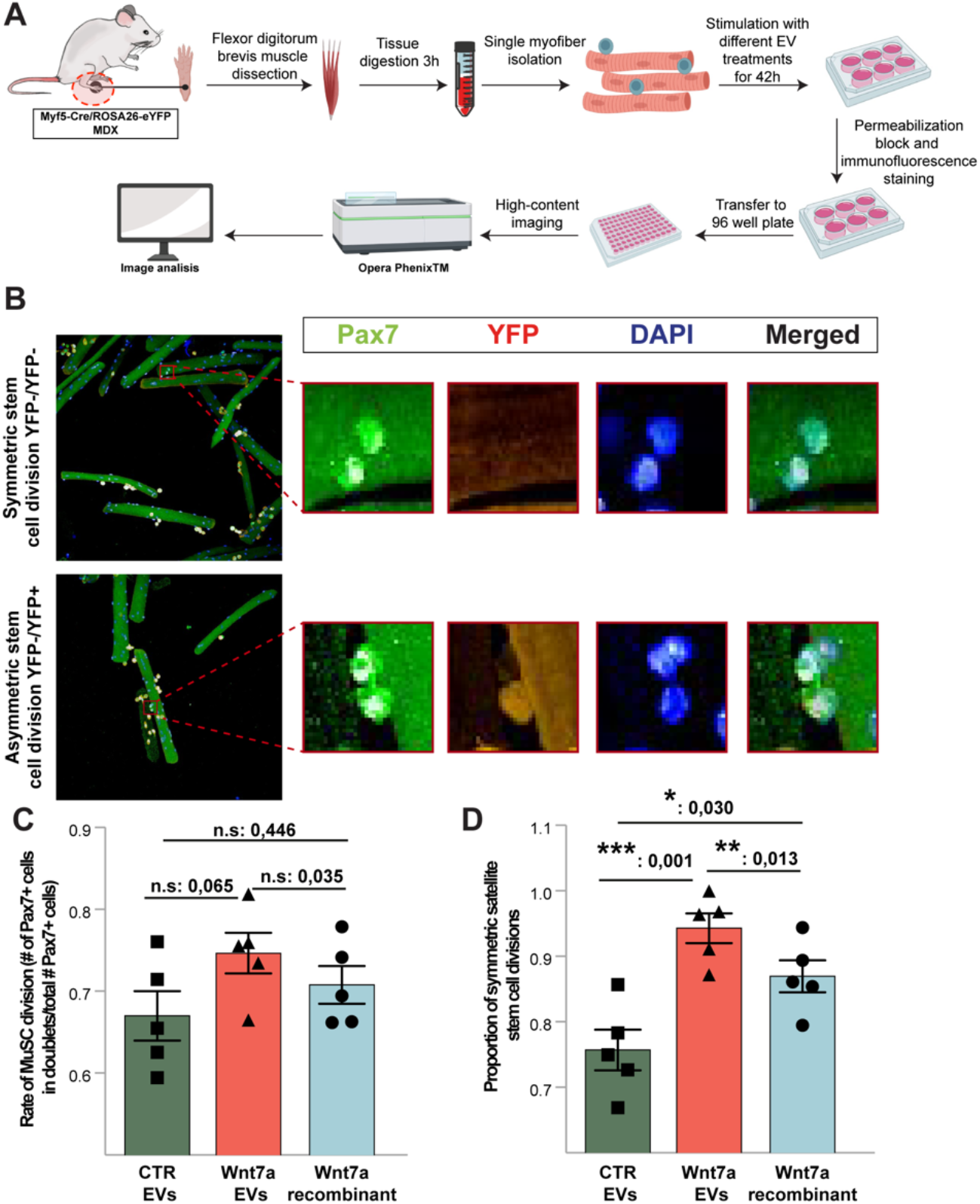
Wnt7a-EVs exhibit enhanced bioactivity. (*A*) Schematic showing the experimental protocol for *ex vivo* EV treatment of single myofibers isolated from FDB muscles, imaging and analysis using high-content screening. (*B*) Representative immunofluorescence images of symmetric (top panel) and asymmetric (bottom panel) satellite stem cell divisions. (*C*) Rate of muscle stem cell division after 42 h in culture is not altered by treatment with either Wnt7a-EVs or with recombinant Wnt7a. (*D*) Symmetric satellite stem cell divisions are significantly increased by Wnt7a-EV treatment. EVs (extracellular vesicles), FDB (Flexor Digitorum fibers). n≥5 replicates, mean ± s.e.m., p values determined by two-sided Student’s test (*p<0.05, **p<0.01, ***p<0.001, n.s.=not significant).

Image analysis was used to quantify the numbers of symmetric and asymmetric satellite stem cell divisions via the expression of eYFP protein (Fig 3B). The rate of muscle stem cell division was not significantly altered by treatment with either EVs or Wnt7a recombinant protein (Fig 3C). However, the proportion of symmetric satellite stem cell expansion was significantly increased by Wnt7a-EV or by rWnt7a treatment (Fig 3D). Interestingly, an equivalent amount of Wnt7a provided on EVs induced significantly more symmetric divisions than rWnt7a suggesting increased bioactivity or bioavailability (Fig 3D). Therefore, we conclude that Wnt7a carried on EVs isolated from transfected primary myocytes displays robust bioactivity.

### Wnt7a-EVs rescues muscle regeneration in *mdx* mice lacking Wnt7a

To assess whether the regeneration deficit observed in *mdx* mice lacking Wnt7a could be rescued by exogenous Wnt7a, we performed tail vein injections using the preparation of Wnt7a-EVs described above. We first evaluated the biodistribution of Wnt7a-EVs following tail vein injection.

We found that following tail vein injection into WT mice, Wnt7a-EVs exhibited uptake by many different organs including heart, spleen, kidney, and liver (Figure S4A). Notably, uptake into skeletal muscle was markedly increased following CTX injury (Figure Sup 4A). Therefore, we asked whether systemically administered Wnt7a-EVs delivered by tail vein injection could ameliorate the dystrophic phenotype in *mdx* mice upon acute injury (Fig. 4A). For these experiments, we made use of the *mdx:Myf5^Cre/+^:Wnt7a^fl/fl^* mice as a sensitized model to assess Wnt7a activity.

**Figure 4:**
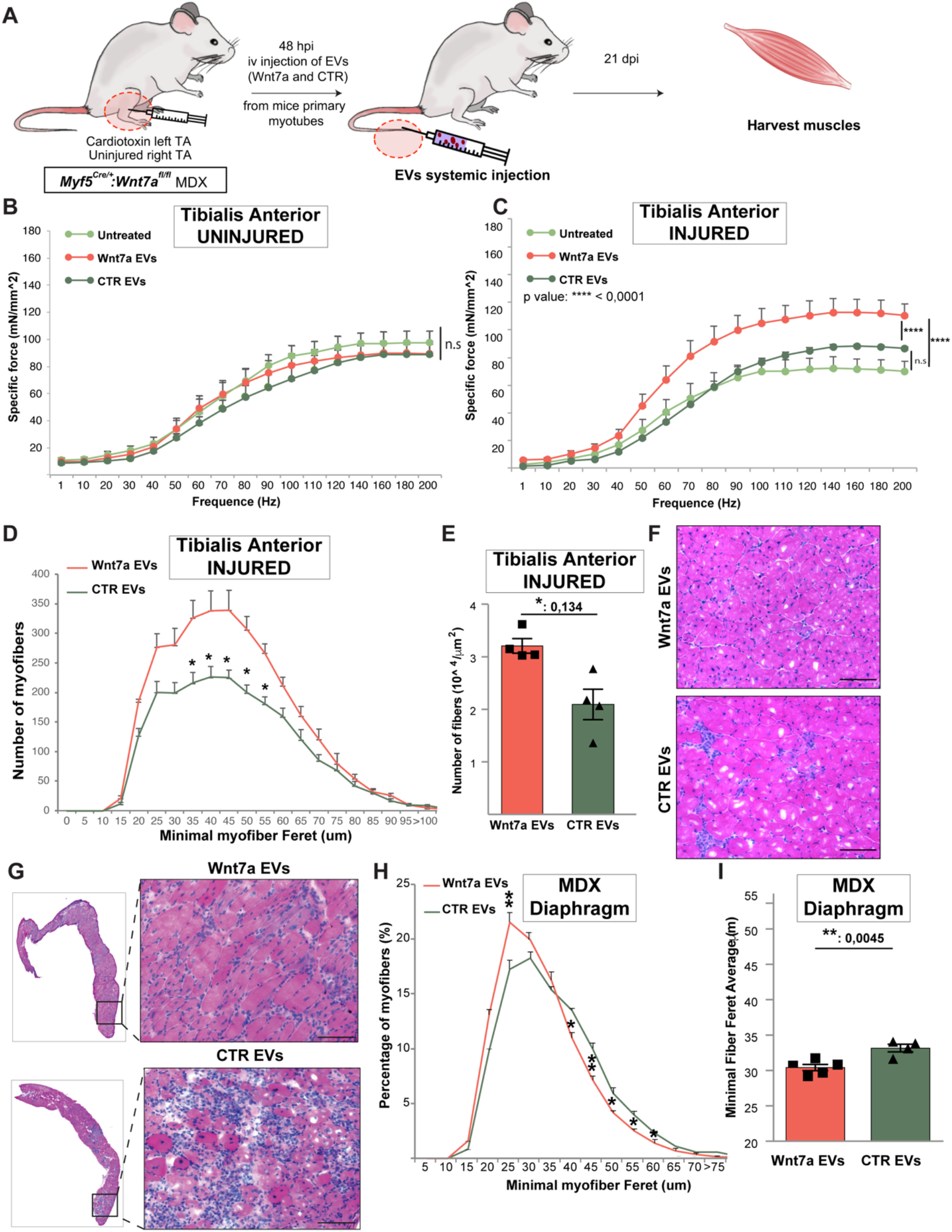
Systemic delivery of Wnt7a-EVs rescues muscle regeneration in *mdx* mice. (*A*) Schematic of experimental design. **(*B*)** Specific Force frequency curve of uninjured *mdx* TA muscle lacking Wnt7a, comparing untreated (light green), Wnt7a-EV (red) injected, or control-EV (dark green) injected. **(*C*)** Specific Force frequency curve of regenerated *mdx* TA muscle lacking Wnt7a, comparing untreated (light green), Wnt7a-EV (red) injected, or control-EV (dark green) injected. **(*D*)** Myofiber caliber distribution of regenerated *mdx* TA muscle lacking Wnt7a, comparing Wnt7a-EV (red) injected, or control-EV (dark green) injected. (*E*) Wnt7a-EV treatment results in increased numbers of myofibers. (*F*) H&E stained sections of regenerated TA muscles after 21 dpi (*G*) Representative H&E stained sections of diaphragms from *mdx* mice lacking Wnt7a intravenously injected with Wnt7a-EVs or control-EVs (*H*) Myofiber caliber distribution of diaphragms from *mdx* mice lacking Wnt7a injected with Wnt7a-EVs or control-EVs (*I*) Minimal fiber feret average in diaphragms of injected mice. TA (Tibialis Anterior), dpi (days post injury. n≥4 mice, mean ± s.e.m., p values determined by two-way ANOVA or two-sided Student’s test (*p<0.05, **p<0.01, ****p<0.0001, n.s.=not significant).

The TA muscles of *mdx* mice lacking Wnt7a were injured by CTX injection and a single dose of Wnt7a-EV or CTR-EV delivered by tail vein injection at 48 h post injury (Figure 4A). The total dose of Wnt7a protein delivered amounted to 700 ng. Muscle force determination and histological assessments were performed at 21 days post injury. In uninjured TA muscles, we observed no change in the force frequency profile (Fig 4B). Remarkably, systemic delivery of Wnt7a-EVs in injured TA muscle had a significant positive effect on the force frequency profile compared to CTR-EVs treatment (Fig. 4C).

Histological characterization of the entire injured TA muscle sections (Fig. S4B) upon systemic treatment with Wnt7a EVs or CTR EVs showed no differences in the distribution of myofiber caliber size (Fig. 4D), average caliber (Fig. S4C) or total cross-sectional area (Fig. S4D). However, systemic treatment with Wnt7a EVs revealed a significant increase in the number of myofibers compared to CTR-EVs consistent with a stimulation of the regenerative response (Fig. 4E). Accordingly, H&E results reflected a better myoregenerative outcome with fewer fibrotic areas after systemic treatment with Wnt7a-EVs relative to CTR-EVs (Fig. 4F). Lastly, no significant difference in Pax7 numbers were found in the treated TA muscle perhaps reflecting the point that a single dose of Wnt7a was delivered 21 days prior to sacrifice allowing a return to homeostasis (Fig. S4E-F).

We lastly asked within the same experiment whether the myoregenerative effect after systemic delivery of Wnt7a-EVs was also stimulating regeneration in the diaphragm. Histological analysis of the diaphragm comparing Wnt7a-EV and CTR-EV treated *mdx:Myf5^Cre/+^:Wnt7a^fl/fl^* mice revealed an overall improvement of the histopathological presentation of the diaphragm condition after treatment Wnt7a-EVs with reduced fibrotic infiltration (Fig. 4G). The distribution of myofiber caliber size (Feret’s dimeter) showed a significant shift towards the formation smaller regenerating myofibers after treatment with Wnt7a-EVs (Fig. 4H,4I and S4H).

All together, these data indicate that Wnt7a-EVS delivered systemically via the circulation positively stimulates regeneration of dystrophic skeletal muscle.

## Discussion

We have discovered that Wnt7a expression in muscle is required for an efficient regenerative response following an acute injury in muscle. Moreover, we observed reduced force generation in muscle of *mdx* mice as well as reduced regenerative capability following an acute injury. The muscle regeneration deficit was rescued by a single tail vein injection of a preparation of Wnt7a-EVs. Therefore, we conclude that intrinsic Wnt7a secretion by myofibers plays an important role in muscle homeostasis and regeneration. Furthermore, our experiments open new therapeutic avenues to develop systemic treatments for Wnt7a therapy for the treatment of DMD.

The lack of Wnt7a expression in WT versus dystrophic muscle has different consequences on the regenerative response. We found that deletion of Wnt7a in WT muscle fibers appeared to delay the myoregenerative response. After acute injury, we observed an increase in Pax7-expressing myogenic cells, no change in the number of myofibers, but a marked reduction in average myofiber caliber, resulting in reduced force generation. By contrast, the deletion of Wnt7a in *mdx* muscle resulted in a severely reduced regenerative response. We observed reduced numbers of Pax7-expressing cells, reduced numbers of myofibers, and a poor overall regenerative response resulting in a decreased ability to generate force. Therefore, we conclude that the relatively efficient regeneration induced by CTX-injection in *mdx* muscle is a consequence of high-level upregulation of Wnt7a in regenerating muscle.

The increase in numbers of Pax7-expressing cells in WT but not in *mdx* muscle following an acute injury suggests that the perturbations of dystrophin-deficient satellite cell function are an important aspect to consider. Dystrophin is highly expressed in newly activated satellite cells where it establishes apical-basal polarity which is necessary for efficient asymmetric divisions, and generation of progenitors (4). Both *mdx* and DMD muscle exhibit a hyperplasia of the muscle stem cell compartment with a 3-5 fold increase in the numbers of Pax7-expressing cells at all stages of the disease (22). It is interesting to speculate that the increase in numbers of WT muscle lacking Wnt7a is a consequence of continued proliferation of progenitors due delayed regeneration. In *mdx* muscle lacking Wnt7a, the Pax7-expressing cells are not generating sufficient progenitors for this response to occur, as evidenced by the marked reduction in the numbers of regenerated myofibers. Further experiments, including single cell RNA-seq would elucidate these phenomena.

Previously, we have shown that endogenous Wnt7a is secreted at high levels *in vivo* on the surface of EVs in regenerating myofibers following an acute injury(7). To generate Wnt7a-EVs for use in our experiments, we expressed Wnt7a in primary myoblasts, then induced differentiation into myotubes to facilitate isolation of Wnt7a-EVs from conditioned using TFF. We found that Wnt7a-EVs exhibited robust bioactivity when we assessed their ability to stimulate symmetric satellite stem cell divisions on cultured myofibers. The enhanced activity of Wnt7a-EVs relative to recombinant Wnt7a may reflect the short half-life of recombinant Wnt7a (23). Alternatively, Wnt7a-EVs may display enhanced bioactivity due to their inherent capacity to facilitate long-range signaling(15–17).

We found that a single injection of Wnt7a-EVs via the tail vein in *mdx* mice lacking Wnt7a rescued their regeneration deficit. Notably, Wnt7a-EV injection resulted in a marked increase in specific force generation compared to mice injected with CTR-EVs. Strikingly, the observed specific generation reaches the same levels as WT muscle following injury. Histological analysis revealed a remarkable improvement in the histopathological appearance of the muscle. Importantly, we confirm that Wnt7a-EVs treatment via intravenous injection represents an efficient systemic treatment to treat the entire body, as histological results from the diaphragm revealed an enhancement of the regenerative response.

Overall, we have demonstrated that Wnt7a is required for efficient muscle regeneration following an acute injury in muscle. Our experiments further reveal a role for endogenous Wnt7a in slowing the progression of dystrophic disease by stimulating regeneration. We have shown that Wnt7a-EVs can be delivered via the circulation to rescue the regeneration phenotype in *mdx* mice lacking Wnt7a. This innovation overcomes the hydrophobicity limitation that characterized Wnt7a free soluble protein, which could only be delivered by intramuscular injection(5). Taken together, our findings suggest that Wnt7a loaded into small EVs represents a potential systemic delivery strategy to stimulate skeletal muscle regeneration in all muscle groups throughout the body toward ameliorating dystrophic progression in Duchenne muscular dystrophy.

## Materials and Methods

### Mice and animal care

All experimental protocols for mice used in this study were approved by the University of Ottawa Animal Care Committee, which is based on the guidelines of the Canadian Council on Animal Care. Food and water were administered *ad libitum*. Muscle regeneration experiments were assessed in 12-week-old male obtained from F2 cross between the offspring of *Myf5-Cre* mice (24) and Wnt7a*^fl/fl^* mice (25) in a C57BL/6 genetic background or C57BL/10ScSn-Dmdmdx/J genetic background. Muscle regeneration experiments were performed as previously described (26) with the following modifications. Mice were anesthetized with isoflurane and CTX injection was performed on a single injection into the TA (50 μl, 10 μM) and muscle regeneration assessed after 21 days. Muscle explants-derived EVs were obtained from C57BL/10ScSn-Dmdmdx/J mice. Briefly, mice were anesthetized with isoflurane and CTX injection was performed on three injections (50 μl, 10 μM each), into the TA, and one on each lobule of the gastrocnemius. Hind limb muscles were harvested after 96h.

### Immunohistochemistry

TA and diaphragm muscle cryosections were rehydrated using PBS, and then fixed with 2% PFA in PBS at room temperature. After washing with PBS, permeabilization with a solution of 0.1% Triton and 0.1 M glycine in PBS was applied for 10 min at room temperature. Mouse on mouse blocking reagent was used at a dilution of 1:40 in blocking solution of 10% goat serum, 1% bovine serum albumin (BSA) and 0.1% Tween 20 in PBS for one hour at room temperature. Primary antibodies were incubated overnight. Nuclei were counterstained with DAPI before mounting in Permafluor. For analysis Z-stack images of cryosections were acquired on an epifluorescence microscope equipped with a motorized stage (Zeiss AxioObserver Z1) with a step size of 0.2 μm to span the cell (25 slices in total) and images were deconvoluted using Zen Software (Zeiss). 3D sum intensity Z-projection was performed with ImageJ software. Laminin staining was analyzed for fiber counting and minimum Feret’s diameter using SMASH (27). Antibodies and dilutions are provided in Table S1.

### *In-situ* TA muscle force measurements

*In-situ* TA muscle function (twitch and tetanic force) was evaluated according to previously published protocols (28, 29). Experimental mice were anesthetized with 2%–5% vaporized isoflurane mixed with O2. The distal TA tendon and TA muscle were exposed, and the mouse was transferred to an *in-situ* mouse apparatus with a temperature-controlled footplate platform. The mouse was positioned on a heated surface to maintain the body and muscle temperature at 37^0^C. The knee was secured to a fixed steel post using surgical sutures and the foot was pinned to the platform to prevent movement from contraction of other muscle groups. The distal tendon of the TA muscle was attached to FT03 force transducers connected to a 79E physiograph (Grass Technologies, Warwick, U.S.A.), which in turn was connected to a KCP13104 data acquisition system (Keithley, U.S.A.). Exposed muscles were kept from drying out using physiological saline solution (118.5 mM NaCl, 4.7 mM KCl, 2.4 mM CaCl2, 3.1 mM MgCl2, 25 mM NaHCO3, 2 mM NaH2PO4, and 5.5 mM D-glucose). Electrical stimulations were applied with two needle electrodes that were placed through the skin just above and below the knee to stimulate the tibial nerve. The electrodes were connected to a Grass S88 stimulator and a Grass SIU5 isolation unit (Grass Technologies). Tetanic contractions were elicited every 100 s with 200 ms trains of 0.3 ms, 5 V (supramaximal voltage) pulses at frequencies ranging from 1 to 200 Hz. Absolute tetanic force was defined as the force generated upon stimulation and calculated as the difference in force at the maximum height of contraction and the force just prior to the stimulation. The specific muscle force was calculated by dividing the absolute muscle force by the muscle cross-sectional area (CSA). Muscle CSA was calculated according to the following equation: CSA = (muscle length in cm / muscle weight in g) * TA muscle density, which was considered to be 1.06 g/cm^3^.

### Conditioned media production for muscle explants derived EVs

For tissue EVs we have standardized a protocol to obtain conditioned media from muscles explants (7). Briefly, four days after injury both hind limbs were harvested and cultured as explants on an exosome-depleted FBS pre-coated dish with high-glucose DMEM (Gibco) and maintained at 37°C in a humidified incubator equilibrated with 5% CO2. After 48h conditioned media was collected for EVs isolation.

### EVs isolation

Conditioned media (20mL) was clarified by sequential centrifugation (300g at RT for 10 min; 2500 g at RT for 10 min and 20,000 g at 4 °C for 20 min). Supernatant was transferred to Flexboy bag (Sartorius) and subjected to tangential flow filtration (TFF) under sterile conditions. Briefly, a KrosFlo Research 2i TFF system (Spectrum Laboratories) coupled to a MidGee Hoop ultrafiltration hollow fiber cartridge (GE Healthcare) 500-KDa MWCO was used. Transmembrane pressure was automatically adjusted at 3 PSI and a shear rate at 3000 s-1. Sample was concentrated up to 10mL and then subjected to continuous diafiltration. Finally, sample was concentrated at 5mL and recovered from the cartridge. Lastly, EVs were pellet down after spinning on an ultrabench centrifuge for 30min at 100,000 g at 4 °C.

### Immunoblot analysis

Immunoblot analysis was performed as described previously (30) with the following modifications. The lysates from EVs were not clarified by centrifugation. The immunoblot transferring was performed onto PVDF membranes. Antibodies and dilutions are provided in Table S1.

### Cell culture

Wnt7a expressing primary myoblasts were obtained upon infection of primary myoblasts from C57BL/10ScSn M1 male mice, previously isolated as Sincennes *et al* (31) described. Briefly, myoblasts passage 4 were seeded in 6-well plate (1.0 X 105 cells/well/4ml media) and infected with lenti-III-Ubc-Wnt7a lentivirus (30ul virus/well) created in our laboratory, empty vector lentivirus as control, containing 6ug/ml Polybrene in 1.5ml culture media (Collagen-coated dishes with HAM F12-X, 10% FBS, 100 U/mL penicillin, 100 U/mL streptomycin and maintained at 37°C in a humidified incubator equilibrated with 5% CO2). Selection was done with Puromycin (2.5 ug/mL) to establish the stable cell line. Six different colonies were picked and Wnt7a expression was verified by immunoblot. Primary myoblasts were cultured on collagen-coated dishes with HAM F12-X, 10% FBS, 100 U/mL penicillin, 100 U/mL streptomycin and maintained at 37°C in a humidified incubator equilibrated with 5% CO2.

### Conditioned media production for cell derived EVs

For EVs derived from Wnt7a expressing primary mytoubes, equal numbers of cells were seeded and let them grow in growth media, previously described. For differentiation, myoblasts were grown up to 80% confluence and growth media was replaced with differentiation medium [HAM F12-X: DMEM (1:1), 5% HS, 100 U/mL penicillin, and 100 U/mL streptomycin] for 3d maintained at 37°C in a humidified incubator equilibrated with 5% CO2. Before differentiation serum was pre-cleared of free of extracellular vesicles as previously stated (32). Then media was collected for EVs isolation following regular protocol.

### Immunogold labeling for EVs

Electron microscopy EV analysis was performed according to the to the protocol previously described by Carrasco-Martinez *et al* (33). Briefly, myotube derived EVs were deposited on colloidal and coated grids and inmmunolabeled with anti-Wnt7a antibody (R&D Systems). First grids were incubated with 0,1% saponine in PBS for 5 min and blocked with 10% fetal bovine serum for 10min. Incubation with the antibody was performed for 45min at room temperature followed by rabbit anti-goat antibodies conjugated to 15nm gold particles (BBI international) for 30min. Finally, grids were washed several times with PBS and double distilled water and negative stained with 1% uranyl acetate in water for 40 seconds. Grids were visualized in a JEM1400 Flash (Jeol, Japan) at 100kV accelerating voltage and a CMOS 4K x4K OneView de Gatan camera (Warrendale, PA, USA). Antibodies and dilutions are provided in Table S1.

### FDB myofiber assay

Single myofibers were isolated from the flexor digitorum brevis (FDB) muscles of 8-week old MDX Myf5-Cre/R26R-eYFP mice according to our published protocol (21). Following dissection, FDB muscles were incubated in DMEM with 2% L-glutamine, 4.5% glucose, and 110 mg/mL sodium pyruvate (GIBCO) containing 0.2% collagenase I (Sigma) for 3 hours at 37^0^C with periodic agitation. Myofibers were isolated using gentle trituration in DMEM+ with 2% L-glutamine, 4.5% glucose, and 110 mg/ml sodium pyruvate (GIBCO) containing 20% FBS (Wisent) with a glass pipet. They were then cultured at 37^0^C in suspension in 24-well plates containing DMEM+ with 2% L-glutamine, 4.5% glucose, and 110 mg/ml sodium pyruvate (GIBCO) containing 20% FBS (Wisent) and 1% chick embryo extract (CEE, Accurate Chemicals), and supplemented with different EVs stimulus, expressing Wnt7a or empty vector. EVs concentration was equal to 10μg/mL of total protein, previously measured with Bratford, accordingly to manufacturer’s instructions. On the case of Wnt7a EVs, 10μg/mL of total protein is equal to 20ng/mL of Wnt7a, based on our quantifications using a standard curve with Wnt7a recombinant protein (Fisher). We have used as a positive control Wnt7a recombinant protein at 100 ng/mL. After 42h in culture, fibers were fixed and stained for Pax7, YFP, and DAPI as previously described (21). Fibers were then transferred to 96-well culture plates (ibidi) and scanned and imaged using the Opera-Phenix high-content screening system, equipped with the Columbus processing and analysis software, which enabled the quantification of satellite cell progenitor and stem cell divisions using an automated approach to reduce bias (21).

### Systemic EVs treatment

All experimental protocols for mice used in this study were approved by the University of Ottawa Animal Care Committee, which is based on the guidelines of the Canadian Council on Animal Care. Food and water were administered *ad libitum*. Muscle regeneration experiments were assessed in 12-week-old male obtained from F2 cross between the offspring of *Myf5-Cre* mice (24) and Wnt7a*^fl/fl^* mice (25) in a C57BL/10ScSn-Dmdmdx/J genetic background. Muscle regeneration experiments were performed as previously described (26) with the following modifications. Mice were anesthetized with isoflurane and CTX injection was performed on a single injection into the TA (50 μl, 10 μM). 48 hours post injury mice were tail vein injected with 25uL saline solution containing myotube derived-EVs, expressing empty vector or Wnt7a and muscle regeneration was assessed after 21 days. Dose was calculated previously in a dose-response assay (data not showed) based on the minimal dose to trigger a regenerative response measured by physiology muscle force assay. Intravenously injected solution of EVs contained 350ug of total protein that equals 430 ng of Wnt7a protein measured by immunoblot, using Wnt7a recombinant protein for the standard curve and diluted in 300uL of saline. Equal volumes and protein concentration were injected for empty vector EVs and Wnt7a EVs.

### Statistical analysis

Experiments were performed with a minimum of three biological replicates and results are presented as the mean ± SEM. Student’s t-test were performed to assess the statistical significance of two-tailed analysis. For multiple comparisons ANOVA test was employed and TUKEY test for post-hoc analysis. *P*-values are indicated as *p ≤ 0.05, **p ≤ 0.01, ***p ≤ 0.001, and *P-*values <0.05 were considered statistically significant. The exact *p* values are provided in the Source Data file.

## Acknowledgments

The authors thank Jennifer Richie for the mice colony management. This study was supported by Defeat Duchenne Canada, the US National Institutes for Health [R01AR044031], the Canadian Institutes for Health Research [FDN-148387; PJT-183804], Ontario Institute for Regenerative Medicine, and the Stem Cell Network. D.D. held a Frederick Banting and Charles Best Canada Graduate Scholarships - Doctoral Award (CGS-D).

---

**Fig. S1.**
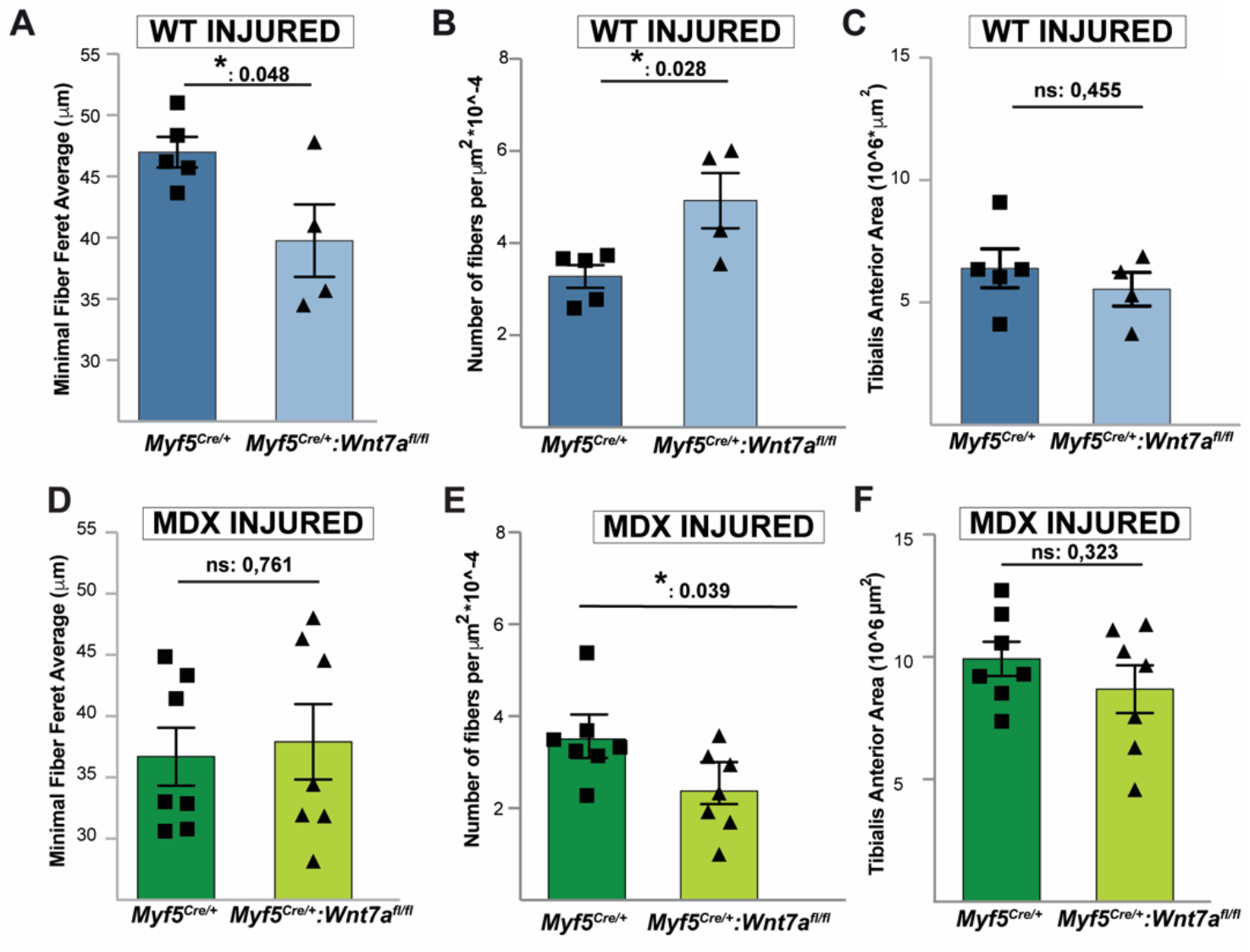
Lack of Wnt7a expression disrupts muscle regeneration. (*A*) Average minimal myofiber Feret at 21 d following regeneration in WT TA muscle with (dark blue) and without (light blue) Wnt7a expression. (*B*) Quantification of myofiber number after regeneration in WT TA muscle with and without Wnt7a expression. (*C*) Total TA area after regeneration in WT TA muscle with and without Wnt7a expression. (*D*) Average minimal myofiber Feret at 21 d following regeneration in *mdx* TA muscle with (dark green) and without (light green) Wnt7a expression. (*E*) Quantification of myofiber number after regeneration in *mdx* TA muscle with and without Wnt7a expression. (*F*) Total TA area after regeneration in *mdx* TA muscle with and without Wnt7a expression. TA (Tibialis Anterior). n≥4 mice, mean ± s.e.m., p values determined by two-sided Student’s t-test (*p<0.05, n.s.=not significant).

**Fig. S2.**
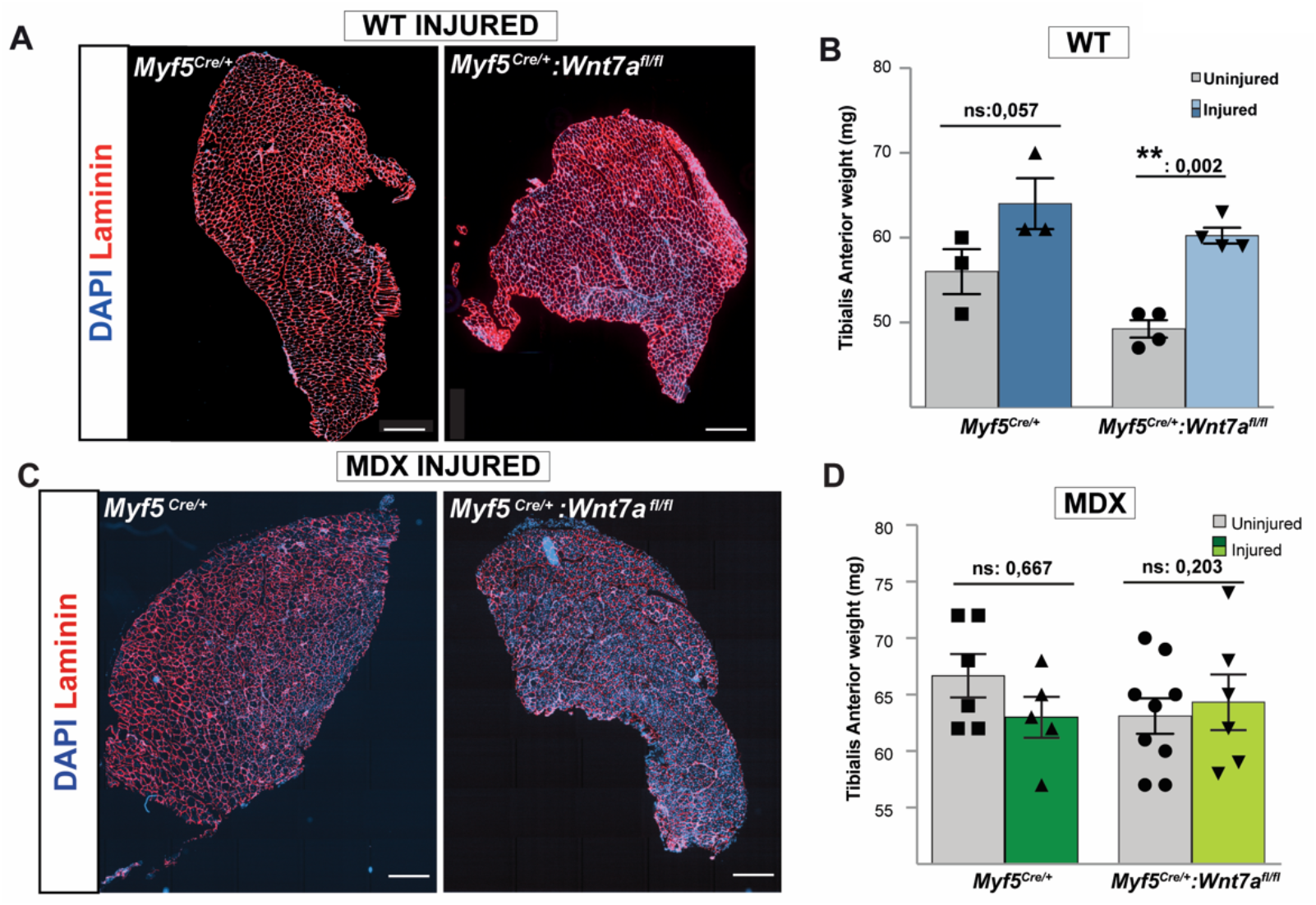
Impaired regeneration of TA muscle lacking Wnt7a. (*A*) Laminin immunostained sections of regenerated WT TA muscle at 21 dpi (*B*) Weight of uninjured and regenerated WT TA muscles (*C*) Laminin immunostained sections of regenerated *mdx* TA muscle at 21 dpi. (*D*) Weight of uninjured and regenerated *mdx* TA muscles. Scale bar 500μm. TA (Tibialis Anterior), Days post injury (dpi), n≥3 mice, mean ± s.e.m., p values determined by two-sided Student’s test (*p<0.05, **p<0.01, n.s.=not significant).

**Fig. S3.**
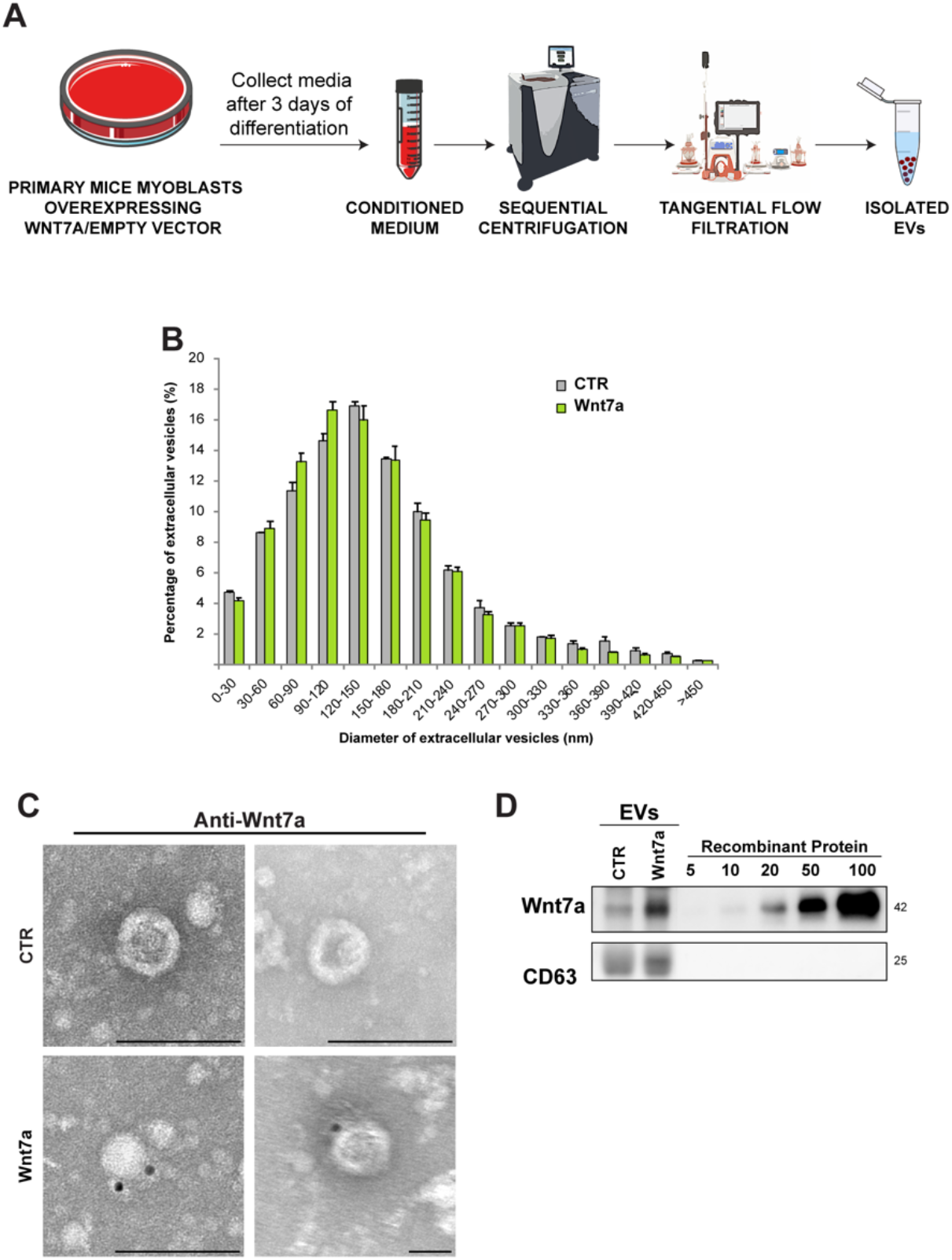
Wnt7a is expressed on the surface of EVs isolated from transfected primary myotubes. (*A*) Experimental protocol used to obtain EVs from myotubes expressing Wnt7a. (*B*) Comparative size distribution analysis of EVs samples after TFF protocol. Experiments and images are representative of n≥3 replicates per condition. (*C*) iTEM of anti-HA immunogold labeling after TFF protocol from Wnt7a expressing myotubes. Scale bar 100nm. (*D*) Quantification of Wnt7a expression on myotube-derived EVs using a standard curve of Wnt7a recombinant protein as a control. CD63 was used as a positive marker for EVs. Tangential Flow Filtration (TFF), Immunogold Transmission Electron Microscopy (iTEM), Extracellular Vesicles (EVs).

**Fig. S4.**
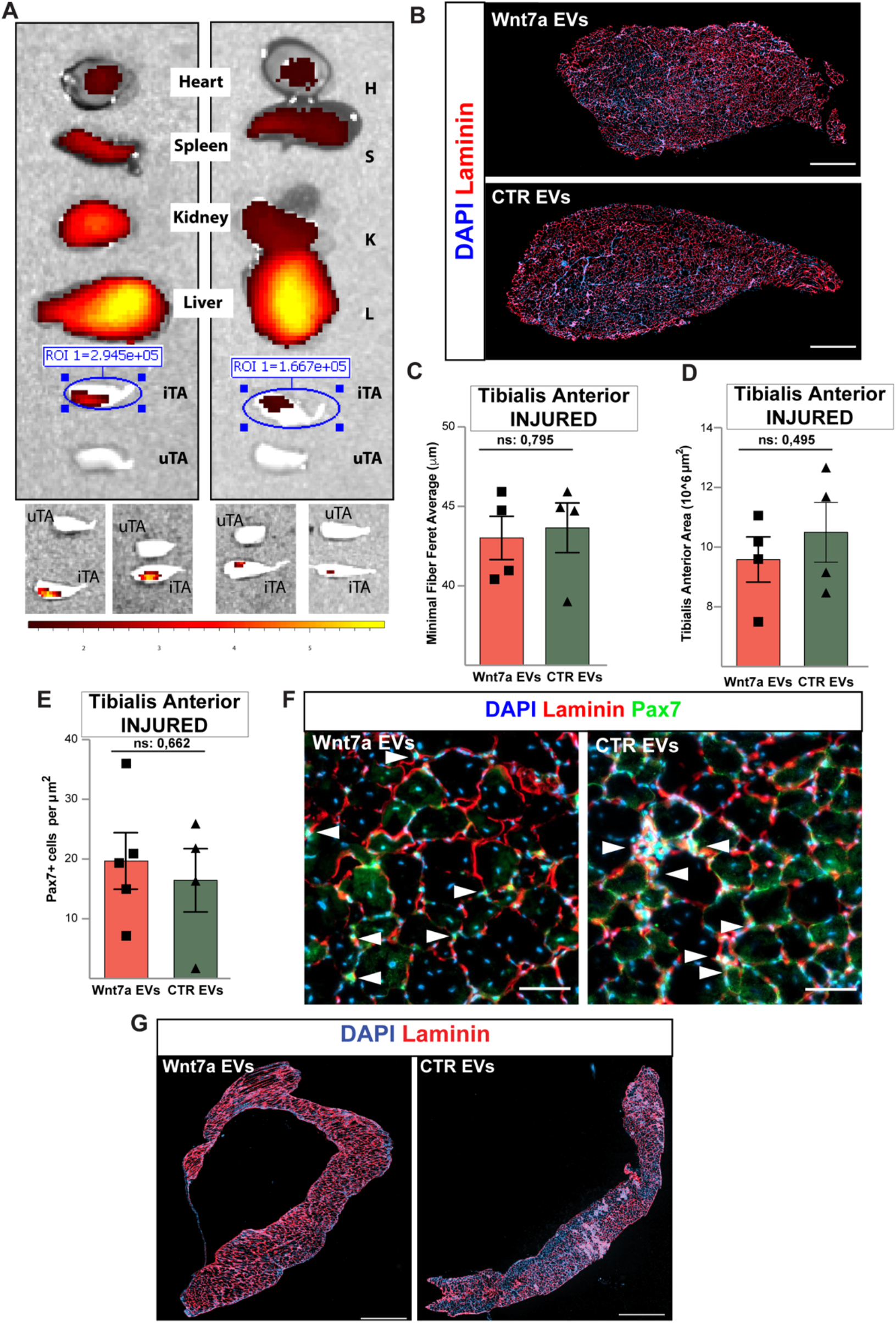
Systemic delivery of Wnt7a-EVs improves acute injury in dystrophic mice. (*A*) Bioluminescence images of labeled EVs uptake in different tissues upon tail vein injection, showing an increased uptake on acute injured Tibialis Anterior (iTA) compared to uninjured Tibialis Anterior (uTA). Bioluminescence quantification in the Region of interest (ROI) (*B*) Laminin immunostaining (red) of TA sections from Wnt7a-EV injected *mdx* mice lacking Wnt7a. Scale bar 500μm. No changes were observed in (*C*) minimal fiber Feret average, (*D*) total TA area, or (*E*) numbers of Pax7-expressing cells. (*F*) Laminin (red) and Pax7 (green) immunostaining of sections of TA muscles from injected mice. Scale bar 50μm. (*G*) Laminin immunostaining (red) of diaphragm sections from Wnt7a-EV injected *mdx* mice lacking Wnt7a. Scale bar 500μm. n≥4 mice, mean ± s.e.m., p values determined by two-sided Student’s test (n.s. not significant).

**Table S1:** Antibodies used in this study.

| **Antibody** | **Application** | **Size** | **Species** | **Dilution** | **Source** | **Secondary Antibody** |
| --- | --- | --- | --- | --- | --- | --- |
| CD63 | Immunoblot | 25KDa | Rabbit | 1:500 | Saint Johns Lab, STJ96890 | Anti-rabbit<br>1:5000 |
| Laminin | Immunofluorescence |  | Rabbit | 1:1000 | Sigma L9393 | Alexa Fluor<br>546<br>(Invitrogen)<br>1:1000 |
| Wnt7a | iTEM Evs |  | Goat | 20ug/mL | R&D Systems AF3008 | 12nm gold<br>Donkey anti<br>Goat. 1:50 |
| Wnt7a | Immunofluorescence |  | Goat | 10ug/mL | R&D Systems AF3008 | Alexa Fluor<br>488<br>(Invitrogen)<br>1:1000 |
| Wnt7a | Immunoblot | 39KDa | Goat | 1:2000 | R&D Systems AF3008 | Anti-goat<br>1:5000 |
| Pax7 | Immunofluorescence |  | Mouse<br>MIgG1 | Undiluted | DSHB PAX7 supernatant | Alexa Fluor<br>488<br>(Invitrogen)<br>1:1000 |
| GFP | Immunofluorescence |  | Chicken | 1/1000 | Abcam ab13970 | Alexa Fluor<br>546<br>(Invitrogen)<br>1:1000 |

